# Dissection of MAPK signaling specificity through protein engineering in a developmental context

**DOI:** 10.1101/115592

**Authors:** Diego L. Wengier, Gregory R. Lampard, Dominique C. Bergmann

## Abstract

Mitogen-activated protein kinases (MAPK) signaling affects many processes, some of which have different outcomes in the same cell. In Arabidopsis, activation of a MAPK cascade consisting of YODA, MKK4/5 and MPK3/6 inhibits early stages of stomatal developmental, but this ability is lost at the latest stage when guard mother cells (GMCs) transition to guard cells (GCs). Rather than downregulating cascade components, stomatal precursors must have a mechanism to prevent late stage inhibition because the same MKKs and MPKs mediate other physiological responses. Here, we artificially activated the MAPK cascade using MKK7, another MKK that can modulate stomatal development, and found that inhibition of stomatal development is still possible in GMCs. This suggests that MKK4/5, but not MKK7, are specifically prevented from inhibiting stomatal development. To identify regions of MKKs responsible for cell-type specific regulation, we used a domain swap approach with MKK7 and a battery of *in vitro* and *in vivo* kinase assays. We found that N-terminal regions of MKK5 and MKK7 establish specific signal-to-output connections like they do in other organisms, but they do so in combination with previously undescribed modules in the C-terminus. One of these modules encodes the GMC-specific regulation of MKK5, that when swapped with MKK7’s, allows MKK5 to mediate robust inhibition of late stomatal development. Because MKK structure is conserved across species, the identification of new MKK specificity modules and signaling rules furthers our understanding of how eukaryotes create specificity in complex biological systems.

## Introduction

MAPKs have strategic roles in signal processing, in mediating stress responses, and in guiding cell fate transitions and development (Keshet and Seger, 2010; Rodriguez et al., 2010; Yang et al., 2013; Chen and Thorner, 2007). MAPK networks consist of a three-tiered cascade whose kinases--MAPK kinase kinase or MKKK, MAPK kinase (Mapk/Erk kinases or MEK in animals, MKK in Arabidopsis) and MAPK (MPK in Arabidopsis) sequentially phosphorylate and activate each other upon signal perception. Downstream effectors may respond to MPK-mediated phosphorylation by changes in protein activity, localization or stability, and many of these alterations ultimately alter transcriptional programs. The MKK level is a bottleneck in many species. In humans, at least 25 MKKKs activate the 7 MEKs, which lie upstream of 14 MAPKs (Keshet and Seger, 2010). In *Arabidopsis thaliana,* more than 60 MKKKs are predicted upstream of 10 MKKs and 23 MPKs (Dóczi et al., 2012). Evidence exists that MKKs can activate more than one MPK, and a given MPK may have more than one upstream MKK (Andreasson and Ellis, 2010). Intuitively, this arrangement could facilitate signal integration, as multiple signals could converge on a single effector. The use of common components, however, could also lead to erroneous cross-activation.

When expressing multiple MAPK network components and responding to multiple sources of information, how do cells generate an appropriate output to a particular signal? One strategy is to make downstream effectors available in only certain cells (Lampard et al., 2008) or under certain conditions (Bao et al., 2004; Brückner et al., 2004; Chou et al., 2004). Alternatively, signaling networks can be insulated through the use of scaffolds, subcellular partitioning of signaling complexes, and by varying signal amplitude or duration (Murphy et al., 2003; Murphy and Blenis, 2006; Raman et al., 2007; Keshet and Seger, 2010; Good et al., 2011; Dóczi et al., 2012). Some MAPK network components encode regions that allow them to establish these connections or localizations. For instance, the animal MEK1/2 uses its proline-rich sequence (PRS) to bind the scaffolds Kinase Suppressor of Ras (KSR) and Mek Partner 1 (MP1). Binding to MP1, in particular, mediates endosomal localization (Schaeffer et al., 1998; Teis et al., 2002; McKay et al., 2009). Several human MEKs and all Arabidopsis MKKs lack a PRS, however, leaving questions about how these smaller MKKs are correctly assembled into restricted MAPK networks.

In Arabidopsis, MAPK signaling has been shown to have a fundamental role in the formation of stomata, the structures in the epidermis of plants that regulate gas exchange (Bergmann et al., 2004; Wang et al., 2007). A systematic study of cell stage-specific responses to MAPK activation revealed that stomatal precursors have mechanisms to limit certain cellular outputs and generate MKK-specific responses (Lampard et al., 2009, 2014). Only four of the 10 MKKs-MKK4, MKK5, MKK7 and MKK9- have any capacity to influence stomatal development during lineage initiation, guard mother cell (GMC) commitment and/or guard cell (GC) formation (for simplicity, only MKK5 and MKK7 are shown in Figure 1). Expression of any of these four constitutively active (CA) MKKs strongly inhibits stomatal formation in early development (Figure 1A, C and D). At the last stage of development, however, MKK4 and MKK5 lose their ability to inhibit stomatal formation (Figure 1B and E). MKK7 and MKK9 activation, in contrast, results in stomatal clustering (Figure 1B and F). Loss of function studies with *MKK4* and *MKK5* indicate that these kinases are endogenously required to limit stomatal production (Wang et al., 2007), but the MKK7/9 significance is unclear as *mkk9* single mutants do not affect stomata and in the recent report of true loss of function mutations in *mkk7* mutants no stomata phenotype was described (Jia et al., 2016).

**Figure 1:**
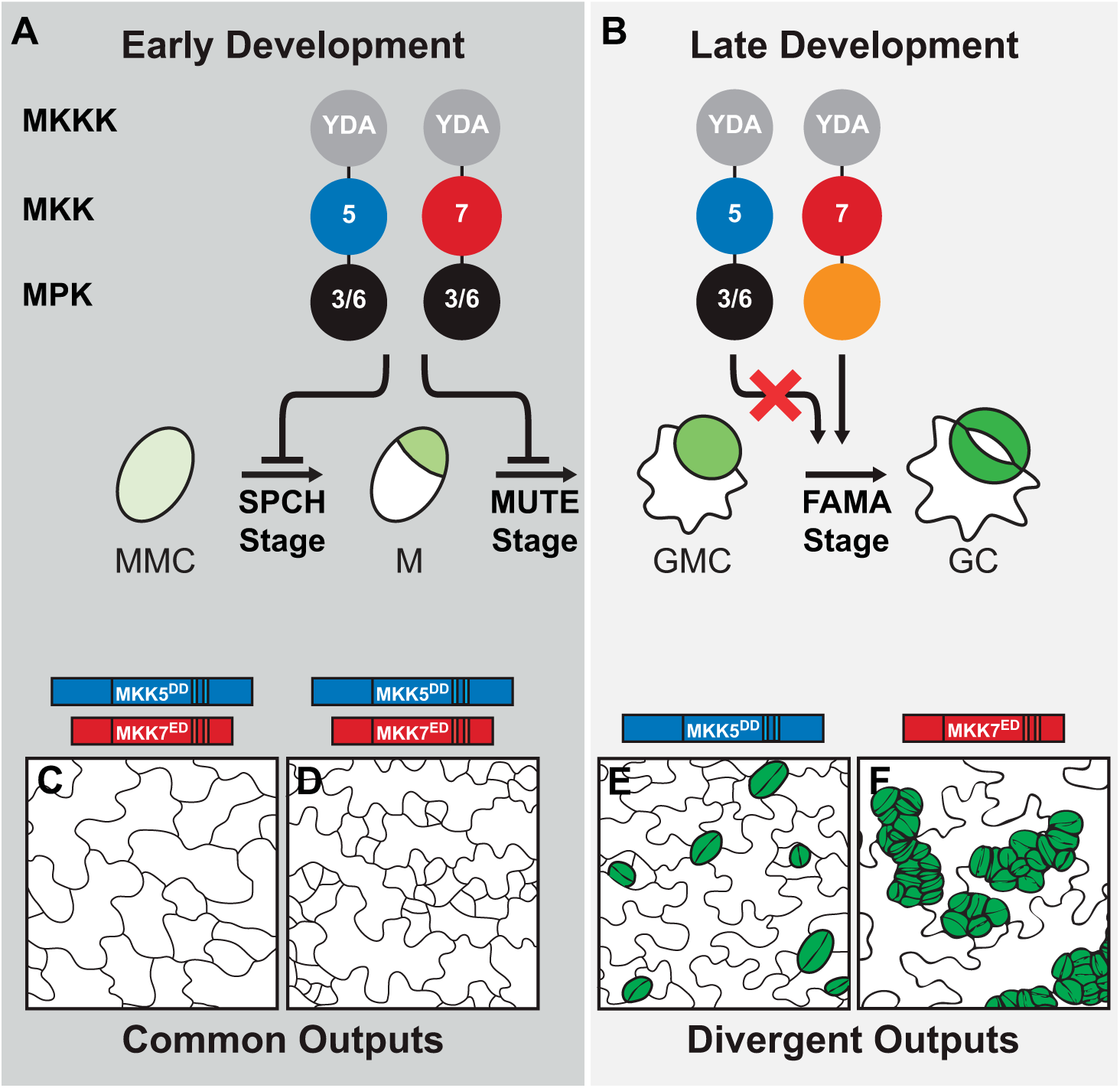
Schematic of stomatal lineage, indicating stages where MKK activation leads to similar and divergent outputs (based on (Lampard et al., 2009)). (A-B) Diagrams of MAPK signaling cascades, with each colored circle representing a different kinase level; circles are labeled with relevant kinase number, with orange circle representing MPK of unknown identity. Constitutive activation of YDA (MKKK), MKK5 or MKK7 inhibits stomatal lineage initiation (A, SPCH and MUTE stages). Late in the lineage (B, FAMA stage) YDA and MKK7, but not MKK5, activation leads to stomatal proliferation via an unidentified MPK. (C-F) Tracings of phenotypes resulting from activation of kinases. In C-D, constitutively active MKK5 (MKK5^DD^) or MKK7 (MKK7^ED^) inhibit initiation (division of meristemoid mother cell (MMC) into meristemoid (M)) and lineage progression (conversion of M into guard mother cell (GMC)). In E-F, MKK5^DD^ has no effect (E, WT numbers and distribution of stomata in green), but MKK7^ED^ induces guard cell (GC) overproliferation and clustering (F). Stages are referred to as SPCH, MUTE and FAMA after the promoters to drive expression of MAPK network components (Lampard et al., 2009). YDA, YODA; 5, MKK5; 7, MKK7; 3/6, MPK3 and MPK6.

MKK4/5’s endogenous role in limiting stomatal production in early stages, but not in late stomatal lineage cells, provides an excellent test case for examining how cell identity may interface with signaling response. We created a quantitative phenotypic analysis pipeline that revealed a previously underappreciated capacity for late stage stomatal lineage cells to be inhibited by MAPK signaling. Then, taking advantage of divergent responses to constitutively activated MKK5 and MKK7, we implemented a protein engineering approach to identify structural domains in MKK5 that are responsible for its stage-specific behaviors. We found that MKK N-terminal regions establish specific signal-to-output connections, much like they do in other organisms (Won et al., 2011), but this requires coordination with previously unexplored regions in the C-terminus. We also found that a minimal domain in the C-terminus encodes the basis for MKK-specific regulation. The location of specificity modules within the plant proteins corresponds to regions in which their human homologues display high sequence diversity, suggesting that these regions may contribute to specificity in many situations. Given the global conservation of MAPK signaling, our findings in the complex multicellular context of plant development may offer insights into general mechanisms of signaling specificity in complex biological systems.

## Materials and Methods

### Plant material and growth conditions

All transgenic lines were generated in the Col-0 background. Seeds were plated on 0.5X Murashige-Skoog media containing 1% agar-agar (Caisson Laboratories, North Logan, UT) and 100 μg/ml Kanamycin (Sigma-Aldrich) or 50 μm/ml Hygromycin B (Life Technologies) when appropriate. Seedlings were grown under a light intensity of 100 μmol photons m^-2^ s^-1^ in a 16:8 photoperiod at 22 ± 1°C. Analyses were performed in 15 day after germination (dag) cotyledons unless stated otherwise. The *mpk6-3* allele was Salk_127507 (Muller et al., 2010). Mutant alleles for *EDR1* (Salk_127158), *MAP3Kδ1* (Salk_048985) and *MAP3Kδ5* (Salk_029929 and Salk_036615) were obtained from the ABRC.

### Multiple sequence alignment and structural analysis

Selected mammalian kinases were aligned using ClustalOmega (Sievers et al., 2011) and structural models are in Supplemental Figure 3B. MKK5 and MKK7 structural predictions were performed with I-Tasser (Zhang, 2008), using MEK1 (1S9J) to assist the prediction. Models were explored with Swiss-PdbViewer v4.1 and fit with the Magic Fit button (Guex and Peitsch, 1997). Structural features were extracted from models and overlaid on the primary sequence (Supplemental Figures 3-5).

### Construction of constitutively active MKK and synthetic chimeras

Domain swap constructs were assembled by fusion PCR from DNA amplicons (blocks) generated with Phusion^®^ High-Fidelity DNA Polymerase following manufacturer’s instructions (New England Biolabs, Ipswich, MA). To generate blocks, MKK5^DD^ and MKK7^ED^ cloned into pENTR without stop codons or other chimeras were used as templates (Lampard et al., 2009). Blocks were designed to contain attL1 and attL2 functional sequences from pENTR to ease the cloning procedure through the Gateway strategy (Supplemental File 1). For domain swaps assembled from two blocks, 5’ blocks contained the M13 forward priming site and attL1 recombination site before the MKK sequence; and 3’ blocks contained the MKK sequence followed by attL2 recombination site and M13 reverse priming site. To facilitate fusion of the blocks, reverse primers for 5’ blocks and forward primers for 3’ blocks were designed as chimeras of the two blocks to be fused, containing at least 15 bases from each block, and were completely complementary to each other. PCR products were gel extracted using QIAquick Gel Extraction Kit (QIAGEN Inc., Valencia, CA) and 1:1 molar ratio mix were used as templates on fusion PCR reactions using M13 forward and reverse primers. Domain swap constructs were gel purified and cloned into pJET 1.2 according to CloneJET PCR Cloning Kit instructions (Thermo Life Sciences, Pittsburgh, PA). For domain swap constructs assembled from 3 blocks, 5’ and 3’ were generated with the same strategy as above, while internal block was amplified with forward and reverse chimeric primers. As domain swaps became more elaborate, first domain swap constructs were used as templates for generating new blocks. Primers, templates and sequences for each domain swap are listed in Supplemental Table 1, A and B.

To build constructs for expression under SPCH and FAMA promoters, 2.5-kb fragments previously described (Ohashi-ito and Bergmann, 2006; Macalister et al., 2007) were first adapted to the Multisite Gateway system. Promoters were shuttled from pENTR to pDONR P4 P1R (Life Technologies, Grand Island, NY) by PCR amplification using promoter shuttling primers (Supplemental File 1) followed by BP recombination performed under manufacturer’s instructions. Promoters flanked by attL4 and attR1 recombination sites (pDONR-promoter) were used in Multisite recombination reactions with domain swap constructs in pJET and R4pGWB440 [destination vector carrying the Gateway cassette flanked by attR4 and attR2 recombination sites, in frame C-terminal fusion to enhanced YFP and kanamycin selection in plants (Nakagawa et al., 2008)]. Recombination reactions were performed in a two-step protocol. First, 1 μl of LR Clonase II was added to 4 μl vector mix (containing 150 ng of pDONR-promoter and 150 ng pJET-domain swap construct) and incubated at 25°C for 5 hours. Then, 150 ng of R4pGWB440 in 4 μl solution were added to the reactions along 1 μl of LR Clonase II. Reactions were incubated for additional 16 hours at 25°C and then stopped after the addition of 1 μl of Proteinase K and incubation for 10 min at 37°C. Constructs were confirmed by sequencing and introduced in Arabidopsis by *Agrobacterium tumefaciens-mediated* transformation.

Mutants *edr1, map3kδ1* and *map3kδ5* were transformed with *FAMA_pro_:MKK5^DD^* in pHGY, previously used in (Lampard et al., 2009). MPK3 and MPK6 clones were provided by Jean Colcombet (INRA Versailles-Grignon, France) (Berriri et al., 2012).

### Scoring phenotypes and data analysis

Seedlings (15 dag) were fixed in 7:1 ethanol:acetic acid, and cleared in Hoyer’s medium. Cotyledons were imaged by differential interference contrast microscopy on a Leica DM2500 microscope at ×20 magnification (0.320 mm^−2^ field of view). One picture per independent transgenic seedling was taken from the distal tip of the cotyledons, within the vascular loop, on the abaxial epidermis. Phenotypes at the FAMA stage were as follows: (1) normal phenotype, only single stomata with tolerance for 1 stomatal cluster per field of view; (2) stomatal inhibition, no stomata present or inhibited precursors coexisted with normal stomata and appeared in at least two independent fields of view per sample; (3) large stomatal cluster, at least two stomatal clusters (4 or more stomata in contact) per field of view; (4) small stomatal cluster, clusters contained 2-3 stomata in contact. When a sample contained a mixed population of clusters, the presence of large clusters defined the classification for this category. MKK7^ED^, clusters were systematically bigger than any chimera and often were delayed in development; to confirm clustering, older epidermis (near apical hydathode or in older plants) was scored. SPCH stage phenotypes were quantified as (1) inhibited (no stomata per field of view) or (2) not inhibited (2 or more stomata per field of view). To enable us to score phenotypes in T1 seedlings that must be grown on antibiotic selection, we grew 35S:YFP lines with the same antibiotic resistance as the MKK variants under the same conditions and scored these as the equivalent of WT controls.

A binomial distribution for phenotypic data was assumed and the percentages of each phenotype were calculated with a confidence of 95%. This analysis is similar to others done on data of comparable nature (Wang et al., 2011). Linear regressions between *in vitro* and *in vivo* data; and between SPCH and FAMA data in Figure 7 and Supplemental Figure 7 were done with Microsoft Excel. To cluster chimeras, hierarchical clustering was performed on phenotypic data at the FAMA stage using the function pvclust in the statistical software R. Percentages of each one of the four phenotypes-Inhibited, Normal, Small Clusters and Large Clusters-were converted to frequencies (e.g. dividing by 100). The distance matrix was obtained by calculating the dissimilarities between all chimeras in their four phenotypes with the Manhattan method. Clustering was performed with the ward.d method and the number of bootstrap replications was 1000. To statistically determine how similar chimeras were within clusters, a Chi-squared test of independence was implemented to compare phenotype distributions. Frequencies were compared to YFP (inactive), MKK5^DD^, MKK7^ED^ (inhibition of stomatal formation and stomatal clustering) and N5-MKK7^ED^ (strong stomatal inhibition). Chimeras that were not statistically different are noted with the same number in Figure 7B. Unnumbered chimeras were statistically different from the rest (p < 0.05).

### Confocal microscopy

Confocal images were collected using a Leica SP5 confocal microscope with excitation/emission spectra of 514/520 to 540 for YFP and 565/580 to 610 for propidium iodide counterstaining. ImageJ (NIH) was used to build Z-stacks from confocal images. To improve resolution of cell outlines, layers were summed rather than averaged. Z-stacks were then split into single channels and only the channel for the cell outlines was conserved, transformed into a grey-scale image and colors were inverted. Bright and contrast were modified to improve image quality. For localization, color channels in Z-stacks were maintained and images were cropped. Localization of MKK-YFP chimeras was investigated in stable (T2-T3) lines with the exception of chimeras that induced stomatal inhibition, where localization was determined in T1s on antibiotic selection.

### Kinase-inactive MPK3 and MPK6 phosphorylation

*In vitro* kinase assays to assess the ability of MKK variants to phosphorylate either kinase inactive (KI) MPK3 or MPK6 were performed as described in (Lampard et al., 2014). KI-MPKs were used to avoid autophosphorylation of the substrates. Band intensity was detected and analyzed using ImageJ (NIH). Each reaction was performed in triplicate. The ratio of phosphorylated KI-MPK/unphosphorylated KI-MPK detected by p42/44 antibody (Cell Signaling, Cat. No. 9102) was used to estimate MKK activity. Background signal corresponds to 1. Mean and standard error was calculated for each sample.

### Yeast two-hybrid assays

Yeast two-hybrid assays was performed with the matchmaker Two-Hybrid System 3 (Clontech) using a modified set of plasmids compatible with Gateway technology and conditions specified by the manufacturer. MKKs and chimeras were cloned as DNA Binding Domain fusions and MPKs were cloned as Activation Domain fusions. Three independent yeast colonies were tested for each pairwise comparison at 1, 1:10 and 1:100 dilutions after incubation of 2, 3 and 4 days in plates containing 1 mM 3 - amino-1/2/4-triazole. Experiments were repeated three times with yeast cultures at OD600 of 1, 2 and 4. Interactions were evaluated as positive if significant growth was detected in 1:100 dilution at day 3.

### Accession Numbers

Arabidopsis Genome Initiative locus identifiers for the genes studied in this work are: SPCH, AT5G53210; FAMA, AT3G24140; MKK5, AT3G21220; MKK6, AT5G56580; MKK7, AT1G18350; MPK3, AT3G45640; MPK6, AT2G43790; EDR1, AT1G08720; MAP3Kδ1, AT1G11850; MAP3Kδ5, AT4G24480.

## Results

### Stomatal development is inhibited by MAPK activation at the FAMA stage

To carefully define the range of phenotypes in our system, we re-analyzed the inhibitory effect of CA-MKK expression in FAMA stage cells (Figure 1B) using more sensitive and quantitative measurements than in our previous studies (Lampard et al., 2009, 2014). For simplicity, we selected one representative MKK from each MKK4/5 and MKK7/9 pair as previous studies showed that MKK4 mirrors MKK5 activity, and MKK7 mirrors MKK9 activity, in every stage of stomatal development (Lampard et al., 2009, 2014). Because MKKs were to be analyzed *in planta,* we selected MKK5 and MKK7, both of which can be easily detected as YFP fusion proteins, thereby providing a control for expression. FAMA promoter (FAMApro) was used to drive the expression of constitutively active MKKs which are made dominantly active by replacing the regulatory S/T residues of the activation loop with phosphomimetic D/E residues (MKK5^DD^ = MKK5^T215D,S221D^ and MKK7^ED^ = MKK7^S193E,S199D^) (Lampard et al., 2009; Popescu et al., 2009). To be able to observe all epidermal phenotypes produced by different MKK expression levels, such as inhibition of stomatal development and stomatal clustering, phenotypes were quantified in cotyledons of independent primary transformants (T1s in Table 1, Figure 2A). We paid special attention to evidence of seedling lethality, a typical result of inhibition of stomatal development.

**Figure 2:**
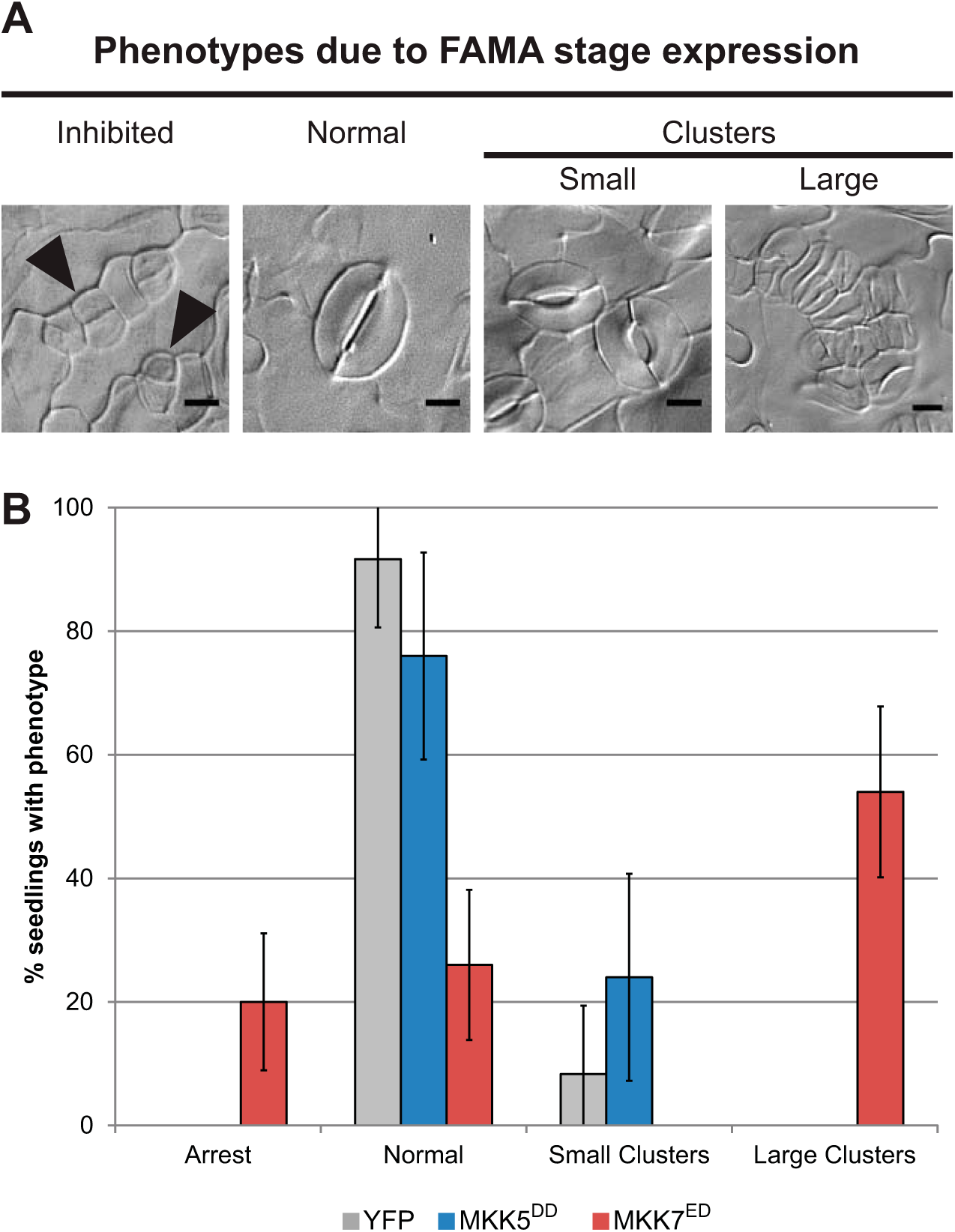
Differences phenotypic output between MKK5 and MKK7. (A) Micrographs of phenotypes resulting from FAMA-stage expression of MKK5^DD^ and MKK7^ED^: inhibited GMCs, where precursors exit the stomatal lineage and do not make GCs (arrowheads); Normal (single stomata comprised of two GCs) or small (2-3) or large (4+) clusters of adjacent stomata resulting from overproliferation of precursors before they become GCs. Scale bars are 10 μm. (B) Quantification of phenotypes, percentage of seedlings showing one of four phenotypes. >19 independent transformants were scored per genotype and stage (Ns reported in Table 1). 35Spro:YFP was used as a negative control (see Methods). Error bars correspond to 95% confidence intervals.

**Table 1:** Quantification of *in vivo* and *in vitro* phenotypes conferred by chimeras.

| Name | Kinase Activity (normalized to MKK5<sup>DD</sup>) |  |  |  | SPCH stage inhibition of lineage initiation |  | FAMA stage phenotypes (as percentage of total T1 seedlings) |  |  |  |  |  |
| --- | --- | --- | --- | --- | --- | --- | --- | --- | --- | --- | --- | --- |
|  | KI-MPK3 |  | KI-MPK6 |  | n | % | n | Inhibited | WT | GC clusters |  |  |
|  | AVG | SE | AVG | SE |  |  |  |  |  | Small | Large | Total |
| YFP control |  |  |  |  |  |  | 24 | 0.00 | 91.67 | 8.33 | 0.00 | 8.33 |
| MKK5<sup>DD</sup> | 34.485 | 9.052 | 38.259* | 2.308 | 69 | 78.26 | 25 | 0.00 | 76.00 | 24.00 | 0.00 | 24.00 |
| MKK7<sup>ED</sup> | 6.836 | 4.808 | 10.867 | 5.726 | 93 | 100.00 | 50 | 20.00 | 26.00 | 0.00 | 54.00 | 54.00 |
| N7-MKK5<sup>DD</sup> | 5.500 | 1.894 | 21.448 | 9.176 | 75 | 26.67 | 125 | 8.80 | 32.00 | 48.80 | 10.40 | 59.20 |
| N5-MKK7<sup>ED</sup> | 15.889 | 7.052 | 35.235* | 1.457 | 22 | 86.36 | 47 | 93.62 | 6.38 | 0.00 | 0.00 | 0.00 |
| MKK5<sup>DD</sup>-C7 | 13.888 | 9.753 | 25.666* | 2.530 | 47 | 87.23 | 44 | 65.91 | 29.55 | 4.55 | 0.00 | 4.55 |
| MKK5<sup>DD</sup>-7A | 45.980 | 22.674 | 26.287 | 4.947 | 114 | 12.28 | 98 | 0.00 | 97.96 | 2.04 | 0.00 | 2.04 |
| MKK5<sup>DD</sup>-7B | 46.746 | 22.671 | 38.718* | 0.750 | 20 | 10.00 | 70 | 50.00 | 50.00 | 0.00 | 0.00 | 0.00 |
| MKK7<sup>ED</sup>-C5 | 3.877 | 2.091 | 0.728 | 0.148 | 43 | 13.95 | 19 | 0.00 | 78.95 | 21.05 | 0.00 | 21.05 |
| MKK7<sup>ED</sup>-CDR5 | 2.451 | 1.242 | 1.303* | 0.312 | 89 | 98.81 | 30 | 96.67 | 3.33 | 0.00 | 0.00 | 0.00 |
| MKK7<sup>ED</sup>-CPR5 | 15.844 | 11.444 | 5.904 | 2.422 | 105 | 0.00 | 36 | 2.78 | 91.67 | 5.56 | 0.00 | 5.56 |
| MKK7<sup>ED</sup>-5A | 15.440 | 12.606 | 5.219 | 4.017 | 125 | 88.00 | 54 | 81.48 | 14.81 | 3.70 | 0.00 | 3.70 |
| MKK7<sup>ED</sup>-5B | 7.238 | 4.545 | 1.317 | 0.200 | 198 | 16.67 | 60 | 0.00 | 48.33 | 40.00 | 11.67 | 51.67 |
| N7-MKK5<sup>DD</sup>-C7 | 3.526 | 1.153 | 2.411 | 0.915 | 91 | 54.95 | 43 | 2.33 | 13.95 | 13.95 | 69.77 | 83.72 |
| N5-MKK7<sup>ED</sup>-C5 | 4.724 | 2.876 | 1.439 | 0.673 | 51 | 72.55 | 46 | 0.00 | 71.74 | 28.26 | 0.00 | 28.26 |
| MPK3, not shown |  |  |  |  | 49 | 0.00 | 50 | 0.00 | 82.00 | 18.00 | 0.00 | 18.00 |
| MPK3<sup>D193G/E197A</sup> |  |  |  |  | 18 | 0.00 | 52 | 0.00 | 80.77 | 19.23 | 0.00 | 19.23 |
| MPK3<sup>T119C</sup>, not shown |  |  |  |  | 43 | 0.00 | 89 | 0.00 | 86.52 | 13.48 | 0.00 | 13.48 |
| MPK6, not shown |  |  |  |  | 122 | 0.00 | 45 | 0.00 | 84.44 | 15.56 | 0.00 | 15.56 |
| MPK6<sup>D218G/E222A</sup> |  |  |  |  | 6 | 100.00 | 27 | 0.00 | 77.78 | 22.22 | 0.00 | 22.22 |
| MPK6<sup>Y144C</sup>, not shown |  |  |  |  | 35 | 0.00 | 21 | 0.00 | 90.48 | 9.52 | 0.00 | 9.52 |

Notes: Kinase activity is determined in triplicates with the exception of those marked with “\*” that were done in duplicates. Averages (AVG) and
Standard Errors (SE) are represented in the table. Phenotypic categories for FAMA stage phenotypes are described in Methods.

Previously, expression of MKK7^ED^ was shown to lead to stomatal hyperproliferation (Lampard et al., 2009), and we could confirm that result: 54% T1s showed large stomatal clusters (Figure 2A and B, Table 1); however, 26% were WT (most of which showed no YFP signal) and 20% had stomatal precursors that failed to complete their development into GCs (Figure 2A and B, Inhibited). This third class died as seedlings. Among MKK5^DD^-YFP transformants, there were no seedling lethals: 76% T1s had a phenotype indistinguishable from controls (Figure 2B, Table 1) and 24% exhibited one to three small clusters (2-3 stomata in contact) per 0.32 mm^2.^. Among control seedlings grown in parallel, ~8% exhibited similar small clusters (Table 1). Phenotype distributions for MKK5^DD^-YFP and MKK7^ED^-YFP were statistically different in Chi-squared test of independence (p << 0.05).

These results indicate that MAPK activation through MKK7^ED^, besides driving stomatal clustering, can also lead to inhibition of stomatal development at the FAMA stage. The failure of MKK5^DD^ to inhibit this stage transition is puzzling, since MKK5 is the endogenous kinase, whereas MKK7 is not normally expressed in FAMA stage cells (Adrian et al., 2015)., Moreover, MKK5^DD^ is an effective inhibitor of earlier stages (Lampard et al., 2009) and *MKK4MKK5* RNAi lines show excess mature GCs (Wang et al., 2007). Also, when compared to MKK7^ED^, MKK5 exhibits stronger interactions with MPK3/6 in Y2H (Supplemental Figure 1) and stronger kinase activity *in vitro* (Supplemental Figure 2). MKK5, therefore appears to be subject to an additional level of *in vivo* regulation that blocks its inhibitory effect, while MKK7 seems to escape this regulation. We reasoned that structural differences between MKK5 and MKK7 could be probed to define the nature and the source of this differential regulation.

### Predicted tertiary structures of MKK5 and MKK7 suggest sources of MKK identity and specificity

We reasoned that the domains most likely to confer the FAMA-stage differential responses would be surface exposed (thus available for interactions with partners) and would exhibit the greatest structural and sequence divergence among MKKs. To facilitate the identification of such regions, we modeled plant MKK folds based on the X-ray crystal structures of human orthologs MEK1 and MEK2 using I-Tasser (Zhang, 2008) (Figure 3) and used structural information from several other mammalian kinases (Hanks and Hunter, 1995; Kannan and Neuwald, 2004; Goldsmith et al., 2007; Knight et al., 2007) to identify conserved features.

**Figure 3:**
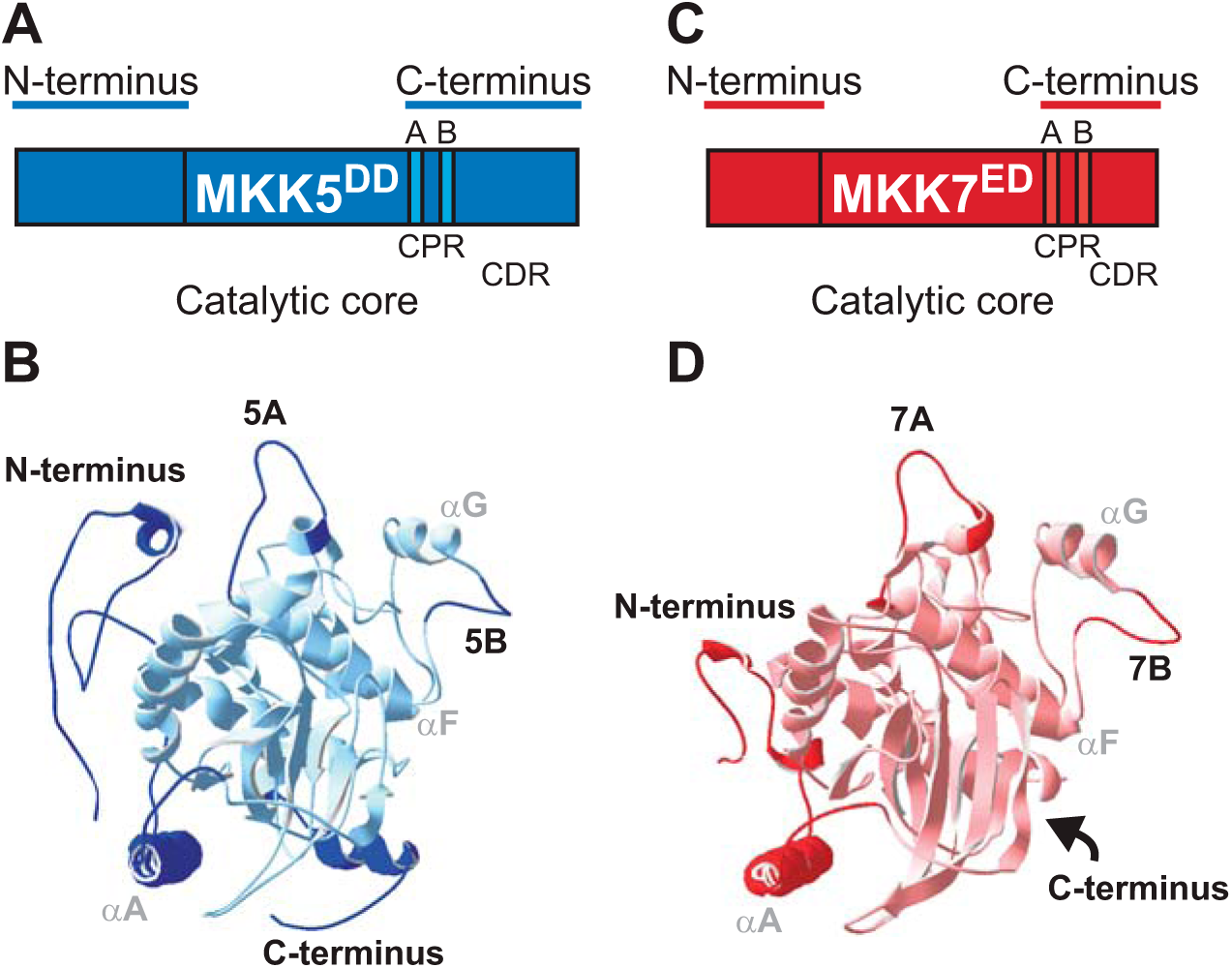
Differences in protein structure between MKK5 and MKK7. (A-D) Schematic and predicted structures of MKK5 (blue) and MKK7 (red). CPR, C-terminal proximal region. CDR, C-terminal Distal Region. (C-D) Four regions important for this study (N-termini, Loop A, Loop B and the C-termini) are bolded. Conserved α helices A, F and G are labeled in grey.

Structural alignment of MKK5 and MKK7 to mammalian kinases confirmed the conservation of the kinase fold (Knight et al., 2007) (Figure 3B and D, with primary sequence in Supplemental Figure 3A). The core catalytic domains of MKK5 and MKK7 are quite similar, but the N- and C-termini are variable. This is similar to MEK1/2, where catalytic domains are similar and well resolved, but the flanking N- and C-terminal extensions and a region containing the PRS are not. In addition, MKK5 possesses, but MKK7 completely lacks, sequences at positions comparable to the C-terminal extension in MEK1/2 (Supplemental Figure 3C). Previously, we showed that this C-terminal extension does not contribute to MKK5 activity or specificity, but that N-termini have an important role in MKKs activities, possibly through the presence of D-docking domains that mediate interactions with downstream MPKs (Lampard et al., 2014).

In addition to the distinct C-terminal distant region (CDR), divergent surface-exposed regions of the plant MKKs include a C-terminal proximal region (CPR) between conserved subdomains VIII and X (Figure 3, Supplemental Figure 3 and 4). The CPR contains two loops. Loop A starts immediately downstream of the YM(S/A)PER sequence, a MKK signature (Rodriguez et al., 2010), and ends before the highly conserved α-helix F. The sequence of loop A is similar between kinases with identical functions and different between kinases with divergent functions in both Arabidopsis and humans (Supplemental Figure 4). For example, loop A in MKK4 and MKK5 is identical, whereas it differs between MKK5 and MKK7. Loop B is downstream of α-helix F and displays a high tolerance for sequence variability (Figure 3, Supplemental Figure 3 and 4). Among CMGC (<u>C</u>yclin-dependent kinases, <u>M</u>APK, <u>G</u>lycogen synthase kinase and <u>C</u>yclin-dependent kinase-like kinase) group kinases, loop B contains an insert that binds interacting proteins (Kannan and Neuwald, 2004), and a different insert in MEK1/2 mediates binding to MAPK scaffolds MP1 and KSR (Schaeffer et al., 1998; Brennan et al., 2011). This region is shorter in plant MKKs (making them resemble human MEK3-MEK7), but the sequence divergence among MKKs is consistent with this loop being a specificity or identity determinant. It is therefore a prime region to target in our dissection of specificity.

### N-termini link specific MKKs to specific phenotypes

Different *in vivo* behaviors of MKK5^DD^ and MKK7^ED^ make it possible to begin to correlate unique sequences and structures with unique functions. Informed by the structural analysis, we made chimeric MKKs based on dividing the MKKs into N-terminal (N), kinase domain (MKK) and C-terminal (C) regions, and further dividing the C domain into CDR and CPR (and within CPR, domain A and B) (Figure 3). To assay the function of these domains, we measured the phenotypes induced by chimeras at the FAMA stage in T1s and compared to those obtained for intact MKK5^DD^ and MKK7^ED^. Expression and subcellular localization of YFP-tagged MKKs was verified by confocal microscopy. We predicted that certain domain combinations could result in non-functional chimeras. Because expression of MKK5^DD^ and non-functional chimeras would give essentially the same phenotype at the FAMA stage (no effect on stomatal development), it was important to discriminate MKK5^DD^-like chimeras from non-functional chimeras. We took two approaches to verify that kinases were still active. First, we measured the intrinsic kinase activity against MPK3/6 in *in vitro* kinase assays (Supplemental Figure 2). Second, we took advantage of the fact that both MKK5^DD^ and MKK7^ED^ drive robust inhibition of stomatal initiation at the SPCH stage (Lampard et al., 2009) to create an *in vivo* assay for kinase activity. We expressed the chimeras at the SPCH stage and quantified the degree of inhibition of stomatal initiation (Table 1).

To characterize the role of the N-terminus in MKK5, we replaced it with the N-terminus of MKK7 (N7-MKK5^dd^) and the chimera was expressed in FAMA stage cells. ~59% of T1 transformants produced stomatal clusters (Figure 4B and G), though clusters were smaller than those generated by MKK7^ED^ In addition, 9% of T1 transformants showed inhibition of stomatal formation (Figure 4G). We also noticed that N7-MKK5^DD^ partially relocalized to mitochondria (Figure 4B) similar to MKK7^ED^ (Lampard et al., 2014). Our *in vitro* and *in vivo* controls for activity both indicated that N7-MKK5^DD^ was less active than intact MKK5^DD^; only ~27% of T1s inhibited stomatal initiation (Figure 4H) and *in vitro* kinase activity was lower, especially towards MPK3 (Figure 4I). This dramatic output alteration (aphenotypic MKK5^DD^ to a weak MKK7^ED^-like behavior) suggests that the N-terminus is more than just a structural/regulatory region required for protein activity. Instead, it appears to channel the MKK towards specific phenotypic outcomes. This specificity behavior resembles that observed in yeast where MKKs involved in other cellular processes were engineered to interact with components of the mating pathway, but were only able to transduce a mating signal when their N-termini were replaced with the N-terminus from Ste7, the mating specific MKK (Won et al., 2011).

**Figure 4:**
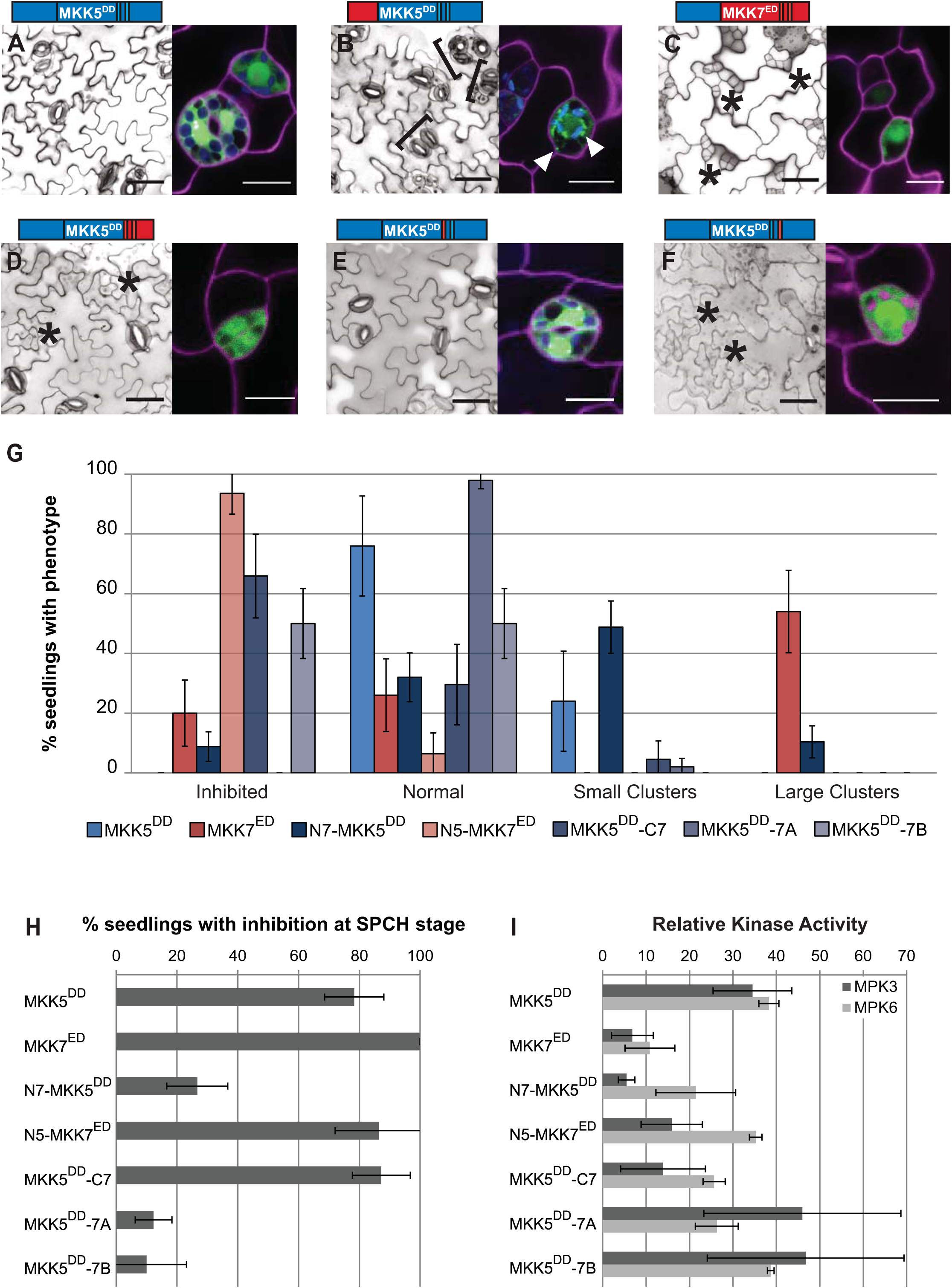
N-termini link MKKs to their phenotypic outputs and the C-terminal Loop B is required for FAMA stage-specific regulation of MKK5. (A-F) Paired micrographs of representative major phenotype (left) and subcellular localization (right), of specific MKK5^DD^- and MKK7^ED^-YFP variants (diagramed above). Black brackets mark clusters and asterisks indicate inhibition. YFP is in green and cell outlines in magenta. For example, in (A) MKK5^DD^ expression results in a WT phenotype and the protein is cytoplasmic and in (B) mitochondrial/cytoplasmic N7-MKK5^DD^ induces stomatal clustering. Scale bars are 50 μm in phenotype images and 10 μm for localization images. (G) Quantification of phenotypes in (A-F). (H) SPCH stage inhibition of lineage initiation. Error bars in (G) and (H) correspond to 95% confidence interval. (I) *In vitro* kinase activity towards kinase inactive MPK3 and MPK6. Kinase assays were performed in triplicates, normalized to unphosphorylated KI-MPK and averaged; error bars represent standard errors.

If the N-terminus enforces MKK specific activities, then replacement of N7 by N5 in MKK7^ED^ should reveal the endogenous response to MKK5 activation. With the N5-MKK7^ED^ chimera we found efficient inhibition of lineage initiation at the SPCH stage and a normal ability to phosphorylate MPK3 and MPK6 *in vitro* (Figure 4H and I). Like MKK5^DD^ the chimera was cytoplasmic localized (compare A and C in Figure 4). Unlike MKK5^DD^, N5-MKK7^ED^ completely inhibited GC production (Figure 4C and G). Thus, with this manipulation, we were finally able to recapitulate the stomatal lineage inhibition phenotype we had expected from MKK5^DD^ based on its ability to inhibit stomatal development at earlier stages (Lampard et al., 2009) and the loss of function stomatal cluster phenotype (Wang et al., 2007).

### Loop B prevents MKK5^DD^ from inhibiting stomata formation at the FAMA stage

Demonstrating that development of FAMA-stage cells could be inhibited, however, raised the question of why intact MKK5^DD^ is unable to do so. We hypothesized that sequences in the MKK5 C-terminus act as negative regulatory regions. To test this idea, we first replaced the entire C-terminal region of MKK5, creating MKK5^DD^-C7. FAMA-stage expression did result in a partially penetrant inhibition of stomatal formation where inhibited precursors coexisted with normal stomata (Figure 4D and G). MKK5^DD^-C7 displayed high activity in SPCH-stage lineage inhibition (Figure 4H), but was less efficient than MKK5^DD^ in *in vitro* kinase assays, particularly towards MPK3 (Figure 4I). Because previously reported MKK5 deletions in the CDR portion of the C-terminus did not significantly change MKK5^DD^ activities (Lampard et al., 2014), we reasoned that putative regulatory regions were located in the CPR.

The largest sequence differences between C5 and C7 reside in loop A and B in the CPR. Substitution of loop A (MKK5^DD^-7A) resulted in a chimera that did not affect stomatal development at the FAMA stage (Figure 4E and G), but substitution of loop B (MKK5^DD^-7B) led to inhibition of stomatal formation at high frequency (Figure 4F and G). This result suggests that MKK5’s loop B is a region that blocks MKK5 from participating in stomatal inhibition at the FAMA stage. Interestingly, SPCH stage activity was markedly reduced for both chimeras (Figure 4H), but *in vitro* activities of MKK5^DD^-7A and MKK5^DD^-7B were at least as high as that of MKK5^DD^ (Figure 4I). These observations suggest that MKK5^DD^-7A and MKK5^DD^-7B are catalytically active kinases but cannot generate appropriate signals *in vivo*.

### Loop B is required for robust MKK7^ED^ activity

If there was truly a discrete domain of MKK5 that was subject to negative regulation, then transferring it to MKK7^ED^ should dampen the stomatal clustering phenotype at the FAMA stage (Figure 5A). We initially swapped the entire C-terminus, and the resulting MKK7^ED^-C5 only produced normal stomata, similarly to MKK5^DD^ (Figure 5B and G). This could suggest that C5 is able to block MKK7 inhibitory function at FAMA stage. However, monitoring other indicators of MKK activity (SPCH stage lineage inhibition and *in vitro* phosphorylation of MPK3 and MPK6) suggested that C7 was essential for overall activity (Figure 5H and I). Thus this phenotype is the result of creating a generally inactive MKK7 mchimera, more than an effect due to the presence of MKK5 regulatory sequences. Thus, we split C5 into CDR5 and CPR5, and determined if we could restore MKK7^ED^ activity *in vitro* and in the SPCH stage. Activity was restored in MKK7^ED^-CDR5: this chimera completely inhibited lineage initiation at the SPCH stage and was indistinguishable from MKK7^ED^ in *in vitro* kinase assays (Figure 5H and I). Rather than decreasing MKK7^ED^ function, however, MKK7^ED^-C**<u>D</u>**R5 had a strikingly stronger inhibitory effect on stomatal development at the FAMA stage than MKK7^ED^ (Figure 5C and G). In fact, it resembled the strong phenotype produced by N5-MKK7^ED^ (Figure 4C). This implies that CDR5, like N5, channels MKK activity to inhibition of stomatal development. In contrast, MKK7^ED^-C**<u>P</u>**R5 was largely inactive at both SPCH and FAMA stages (Figure 5D and G), and in *in vitro* phosphorylation assays against MPK3 and MPK6 (Figure 5H and I), indicating that CPR7 is necessary for MKK7^ED^ catalytic activity.

**Figure 5:**
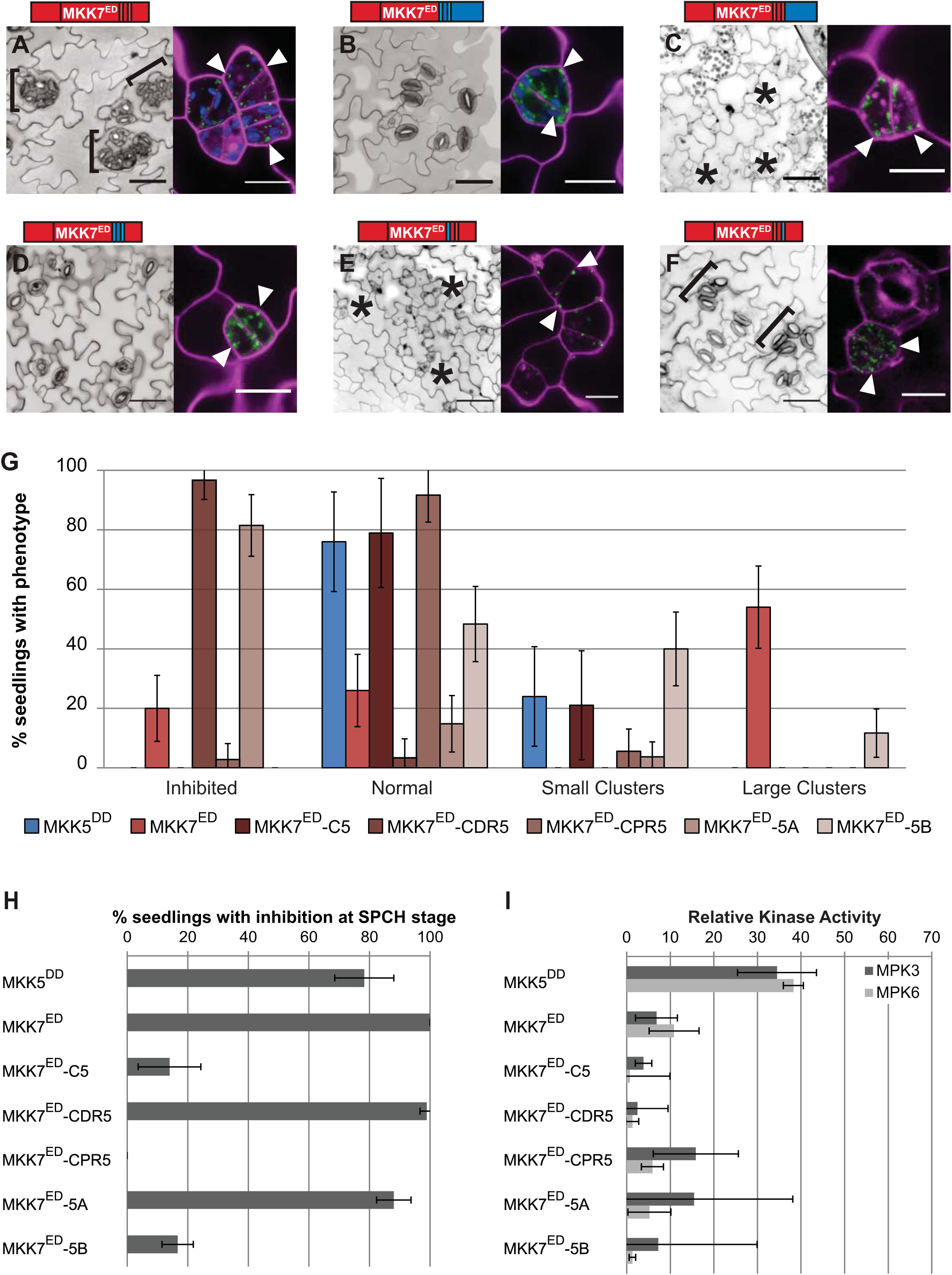
Loops A and B are required for specific and robust FAMA stage MKK7 activities. (A-F) Paired micrographs of representative major phenotype (left) and subcellular localization (right), of specific MKK7^ED^-YFP variants diagramed above. Black brackets mark clusters and asterisks indicate inhibition. YFP is in green and cell outlines in magenta; white arrowheads point to mitochondrial localization. For example, in (A) MKK7^ED^ induces stomatal clustering (brackets) and is mitochondrial localized (white arrowheads). Scale bars are 50 μm for phenotype images and 10 μm for localization images. (G) Quantification of phenotypes in (A-F). (H) SPCH stage inhibition of lineage initiation. Error bars in (G) and (H) correspond to 95% confidence interval. (I) *In vitro* kinase activity towards kinase inactive MPK3 and MPK6. Kinase assays were performed in triplicates, normalized to unphosphorylated KI-MPK and averaged; error bars represent standard errors.

So far, the domains from MKK5 that dampened MKK7^ED^ activity at the FAMA stage also decreased the activity of the chimeras *in vitro* and at the SPCH stage (MKK7^ED^-C5 and MKK7^ED^-CPR5). From these results, it appears that the CPR region is important for MKK7^ED^ catalytic activity and thus we created smaller domain swaps (loops A and B) to attempt to transfer negative regulatory sequences from MKK5 into MKK7^ED^ without affecting kinase functionality. MKK7^ED^-5A and MKK7^ED^-5B were active *in vivo* (inhibited lineage initiation at the SPCH stage, Figure 5H) and *in vitro* (phosphorylated MPK3/6, Figure 5I), although to different degrees. At the FAMA stage, MKK7^ED^-5A inhibited stomatal formation to a greater extent than MKK7^ED^ (Figure 5E and G). This behavior is similar to N5-MKK7^ED^ (Figure 4C) and MKK7^ED^-CDR5 (Figure 5C), suggesting that N5, loop 5A and CDR5 restrict MKK7^ED^ activity to inhibition of stomatal development. In contrast, MKK7^ED^-5B’s ability to cause stomatal clustering and inhibition of stomatal development at the FAMA stage was markedly reduced when compared to MKK7^ED^ (Figure 5G). Because MKK7^ED^-5B also showed reduced activities in other indicators of MKK activity (Figure H and I), we concluded that the negative regulation of MKK5^DD^ is restricted to loop 5B but cannot be transferred without affecting MKK7^ED^ catalytic activity. Nevertheless, the same results highlight that loop 7B is required for robust MKK7^ED^ activity.

### Swapping domains allows specificity to be changed in MKK5^DD^ and MKK7^ED^

Our results show that the N-terminus, CPR region (loops A and B) and CDR region modulate MKK activity. We showed that N7 and CPR7 are necessary for MKK7^ED^-mediated GC clustering at the FAMA stage, but when CDR5 is incorporated into MKK7^ED^, GC production is inhibited. If our “wiring diagram” for specificity is correct, then a chimera that contains the GC promoting domains from MKK7 but not the inhibitory CDR5 (i.e., N7-MKK5^DD^-C7) should mimic MKK7^ED^. Indeed, when we constructed N7-MKK5^DD^-C7, it resembled MKK7^ED^ both qualitatively and quantitatively (Figure 6A and C). Likewise, N5-MKK7^ED^-C5 should match MKK5^DD^ activities, and it does *in planta* (Figure 6B and C). Interestingly, robust rewiring *in vivo* (Figure 6C and D) appears to be uncoupled from kinase activity *in vitro,* as both rewired proteins were much less capable of phosphorylating MPK3 and MPK6 than MKK5^DD^ and MKK7^ED^ (Figure 6E). One interpretation of these swaps is that specificity lies only outside of the kinase domain. If this were true, then we should be able to generate a chimera that resembles N7-MKK5^DD^-C7 and MKK7^ED^ using the kinase domain from another MKK. We selected the kinase domain of MKK6 that can also phosphorylate MPK3 and MPK6 *in vitro* (Popescu et al., 2009), and created N7-MKK6^DD^-C7. Expression of N7-MKK6^DD^-C7 at the FAMA stage, however, did not produce any noticeable phenotype (Supplemental Figure 6). This suggests that although kinase domains in MKK5 and MKK7 are not differential, they still contain stomatal fate-enabling regions.

**Figure 6:**
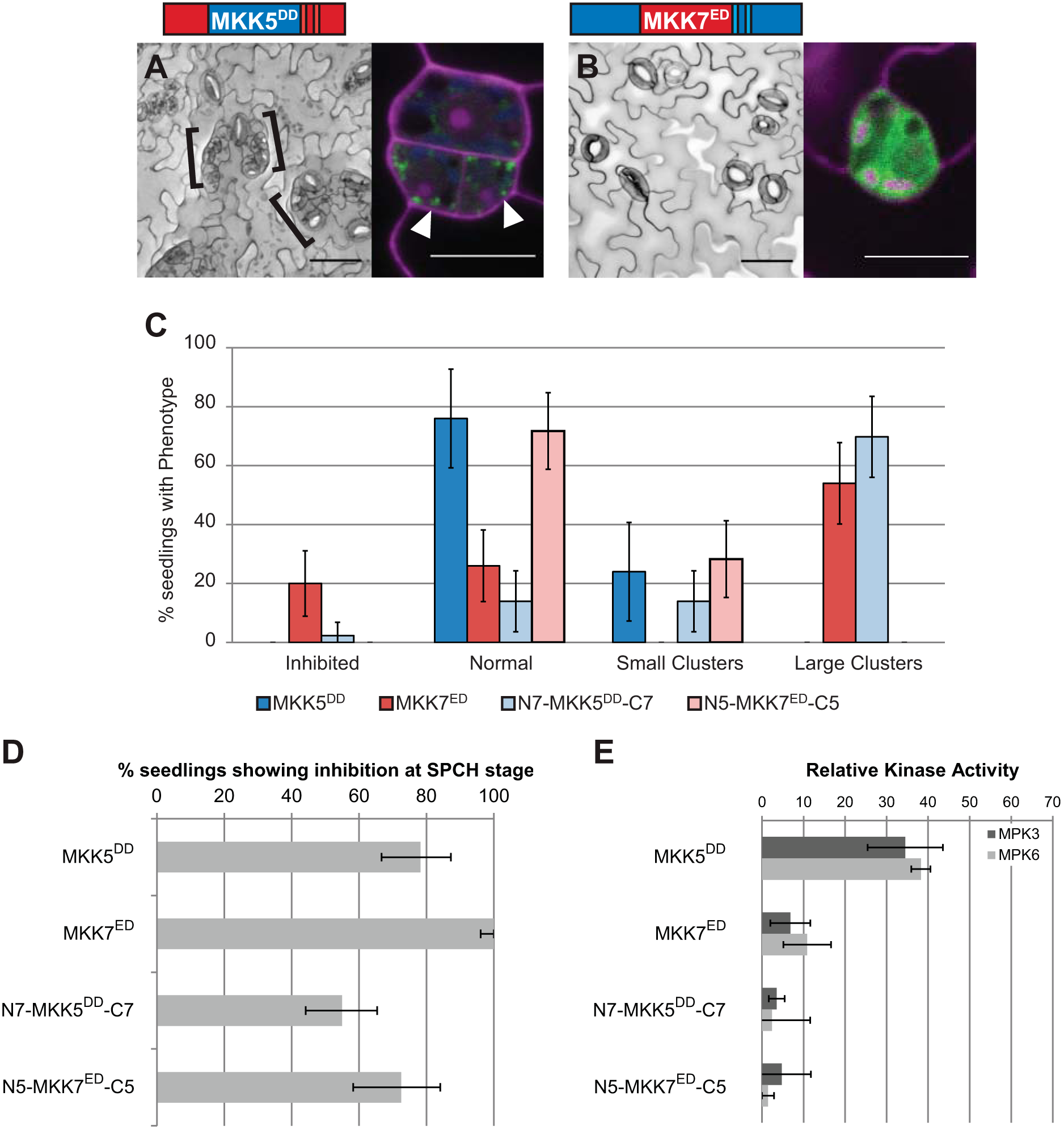
MKK5 and MKK7 activity and localization in the stomatal lineage can be reciprocally rewired. (A-B) paired micrographs of representative major phenotype (left) and subcellular localization (right), specific MKK-YFP variants diagramed above. Black brackets mark clusters; YFP is in green and cell outlines in magenta; white arrowheads point to mitochondrial localization. (A) N7-MKK5^DD^-C7 mimics MKK7^ED^ in that it produces clusters and can localize to mitochondria. (B) N5-MKK7^ED^-C5 mimics MKK5^DD^ in that it has a WT phenotype and localizes in the cytoplasm. Scale bars are 50 μm in phenotype images and 10 μm for localization images. (C) Quantification of phenotypes (A-B). (D) SPCH stage inhibition of lineage initiation. Error bars in (C) and (D) correspond to 95% confidence interval. (E) *In vitro* kinase activity towards kinase inactive MPK3 and MPK6. Kinase assays were performed in triplicates, normalized to unphosphorylated KI-MPK and averaged; error bars represent standard errors.

### Comprehensive analysis of chimeras reveals functions of MKK domains

We repeatedly observed that the ability of native and chimeric MKKs to phosphorylate their targets *in vitro* does not predict their activities *in vivo.* In fact, when chimera data are considered together, *in vitro* versus *in vivo* data have no statistical correlation (Supplemental Figure 7). In contrast, when only *in vivo* data were compared, activities in SPCH and FAMA stages were positively correlated (Figure 7A). Interestingly, native and chimeric MKKs were distributed in two subpopulations. MKKs closer to the regression line promoted stomatal clustering (red dots) or inhibited stomatal formation (black dots). MKKs further from the regression line had no effect in stomatal development at the FAMA stage (blue dots), but had a broad range of activities at the SPCH stage (shaded area in Figure 7A). This behavior might be reflecting the additional regulation that some of the MKKs showed at the FAMA stage.

**Figure 7:**
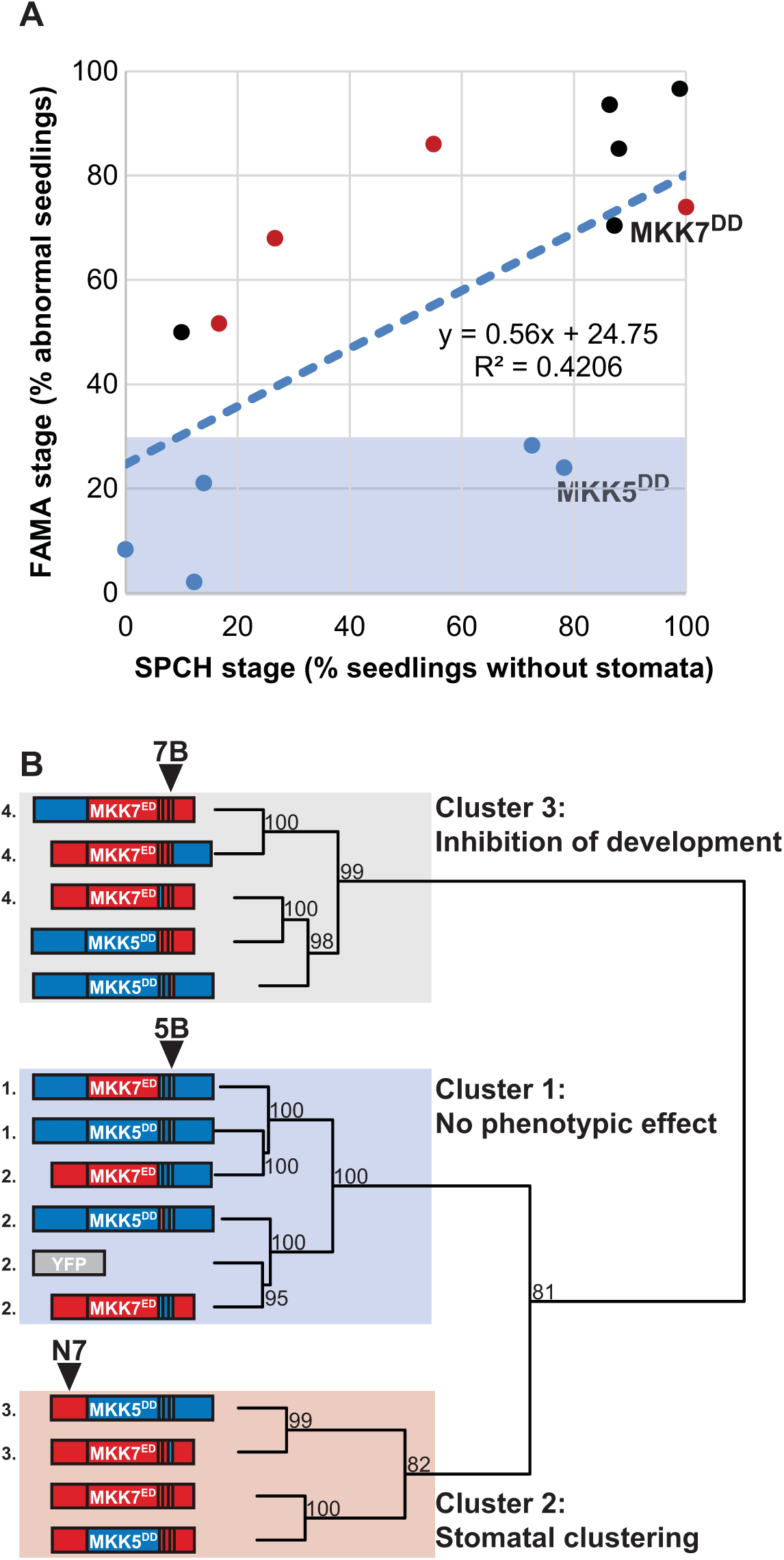
MKK5 defines a cluster of chimeras with low activity at FAMA stage, whereas clustering and inhibiting chimeras group in two independent clusters. (A) SPCH and FAMA stage activities show a weak positive correlation. Shaded area corresponds to activities at the FAMA stage lower than 30%. Data points are colored according to the cluster they belong to in (B). MKK activities at the FAMA stage are separated into three clusters (B): cluster 1 contains MKKs that show no phenotypic effect and contain loop 5B; cluster 2 contains MKKs that lead to stomatal clustering and contain N7; and, cluster 3 contains MKKs that lead to inhibition of stomatal development and contain loop 7B. Numbers next to nodes correspond to approximately unbiased *p*-values with bootstrap replications set to 1000. Chimeras not significantly different in Chi-squared tests if independence are indicated with an identical number (p > 0.05).

We reasoned that MKKs subject to the same regulation would share structural similarities. To test this hypothesis and generate an overall picture of the relationship between MKK structural domains and *in vivo* functions, we clustered 15 native and chimeric MKKs and controls according to their quantitative phenotype data at the FAMA stage (clustering detailed in methods). Constructs robustly fell into three clusters (Figure 7B): Cluster 1, no phenotypic effect; Cluster 2, induces stomatal proliferation; and Cluster 3, inhibits stomatal formation (the presumed endogenous role for MKK5). Within clusters, however, not all MKKs were identical. We performed sequential tests of independence to determine how similar the distribution of phenotype frequencies was between chimeras in each cluster. Cluster 1 was composed of MKKs similar to MKK5^DD^ (group 1) and MKKs similar to inactive YFP (group 2). Cluster 2 was statistically separated into weak MKK7^ED^-like chimeras (group 3) and two MKKs that induced strong clustering, yet were different from each other. Cluster 3 was statistically separated into strong inhibitors of stomatal formation (group 4) and weak inhibitors (different from each other).

To summarize, when analyzing native and chimeric MKK structures across Clusters, we see that loops A and B have discrete functions in selecting MKK-specific outputs and in kinase activity *in vivo.* Loop A can be thought of as a “channel selector”, that, together with N-terminus and CDR, selects between the normal role of arresting stomatal progression and the artificial role of promoting stomatal clustering. Loop B is a “volume control” with the 7B version increasing, and 5B decreasing, the phenotypes specified by the other domains of the MKKs

### MPK6 mediates GC inhibition, but like MKK5, is prevented from doing so at the FAMA stage

Previous loss- and gain-of-function experiments placed MPK3 and MPK6 downstream of an activated MKK4/5 homologue (NtMEK2), suppressing stomatal formation (Wang et al., 2007) in the early stages of the stomatal lineage, but this assay could not address the potential for MPK3 and MPK6 to mediate FAMA-stage activities. Our chimeras that drive stomatal inhibition at the FAMA-stage, however, could be used to see whether either mediated such late stage inhibition. We used N5-MKK7^ED^ which, in WT led to complete inhibition of stomatal development (Fig 8C, Table 1). When expressed in the loss of function *mpk6-3* background, N5-MKK7^ED^ failed to promote complete inhibition in 19 independent T1s (Figure 8D) indicating that MPK6 is likely downstream. This led us to the question of whether MPK6, like MKK5^dd^, would also be actively inhibited from effecting fate at the FAMA stage. To test this, we created a constitutively active MPK6 (MPK6^DE^) (Berriri et al., 2012) and tested its ability to suppress stomatal formation. Expression of MPK6^DE^ (but not MPK3^DE^, Figure 8A, Table 1), inhibited stomatal progression at the SPCH stage, indicating that M8PK6^DE^ is active in this assay. When expressed at the FAMA stage, however, MPK6^DE^ did not affect stomatal development (Figure 8B, Table 1), a phenotypic output remarkably similar to that of MKK5^DD^ (early, but not late, inhibition). We hypothesized that MKK5 and MPK6 normally repress stomatal development, but are actively prevented from having this effect at the FAMA stage.

**Figure 8:**
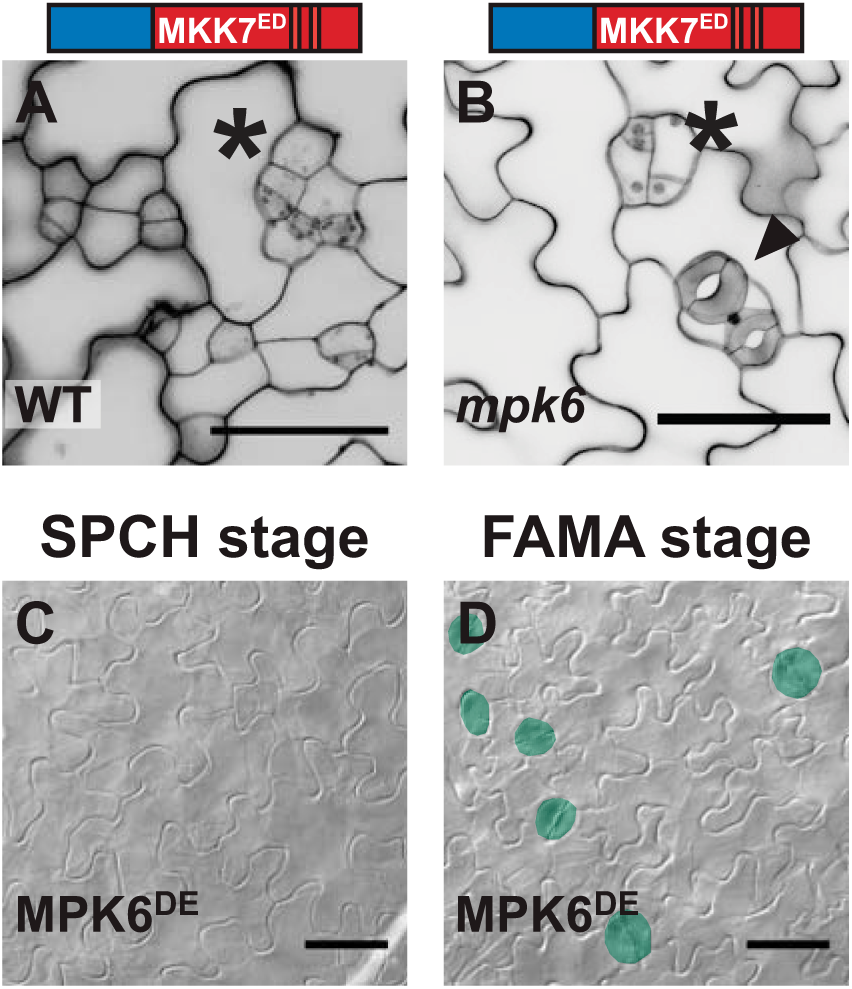
MPK6 inhibits stomatal development at the FAMA stage in a MKK-specific manner. (A and B) Micrographs indicating the requirement for MPK6 as a downstream factor in N5-MKK7^ED^-mediated FAMA-stage inhibition (asterisks). In *mpk6* mutants (B), stomata (arrowhead) coexist with precursors that exit the lineage. (C-and D) DIC micrographs showing results of expression of MPK6^DE^ at SPCH and FAMA stages, stomata are highlighted in green. MPK3^DE^ does not affect development at any stage (Table 1). Scale bars in are 50 μm.

## Discussion

In multicellular organisms, coordinated development requires constant communication between cells and the evaluation of environmental conditions. All this information is integrated to decide from a spectrum of possible outputs, and the spectrum is frequently limited by a cell’s identity. In previous, more superficial studies, FAMA stage cells appeared to lose the ability to inhibit stomatal development upon MAPK activation (Lampard et al., 2009). Here we show that these cells do not lack the capacity to be inhibited, but rather that MKK5 (and possibly MPK6) may be actively prevented from participating in this cellular outcome. Structural analysis and engineered chimeras revealed that this regulation and other specific responses rely on distinct MKK domains.

We found that Arabidopsis MKKs behave as modular proteins with four discrete regions: N-terminus, CDR and two loops (A and B) in the CPR. N-termini contribute to subcellular localization (Figure 4 and in (Lampard et al., 2014)), to phenotypic output (Figure 4) and may mediate interactions with downstream MPKs through their docking domains. In particular, we hypothesize that N7 has the ability to bind different types of MPK. Throughout development, MKK7 inhibits stomatal development by recruiting MPK3/6, but a yet unknown proliferative MPK mediates stomatal clustering at the FAMA stage (Figure 9A). In the C-termini, Arabidopsis MKKs 1, 2, 4, 5 and 6 contain an extension that could be equivalent to the MKKK-interacting domain for versatile docking (DVD) in human MEKs 1, 3, 4, 6 and 7 (Supplemental Figure 4) (Takekawa et al., 2005). Arabidopsis MKK7, 8, 9, and 10, however, lack this domain, making it unclear how they engage the appropriate MKKK. In fact, the addition of CDR5 to MKK7^ED^ restricted this kinase’s activity to an inhibitory output (Figure 5C and G), suggesting that CDR5 interferes with MKK7^ED^ interactions. Upstream of the CDR, Loop A and B are two surface-exposed modules in the CPR that may contribute to establishing interactions with other network components. In our experiments, swapping loop A in MKK7^ED^ restricted its phenotypic output such that MKK7^ED^-5A only inhibited stomata formation (Figure 5E, G). We propose then, that loop A promotes certain MKK-MPK interactions or, alternatively, restricts how MPKs contact MKKs. This hypothesis is supported by sequence similarities between human MEKs that share the same downstream MAPKs (Keshet and Seger, 2010). For example, ERK kinases MEK1 and MEK2 have identical loop As, and p38 kinases MEK3 and MEK6 differ at only one site (Supplemental Figure 4). Interestingly, human MEK7, which can phosphorylate both JNK and p38, shares some residues with MEK3 and MEK6 and others with the JNK kinase MEK4.

The function of loop B seems to be associated to MKK-specific regulation. Our data shows that loop B is required for robust MKK7^ED^ activity, but it prevents MKK5^DD^-mediated inhibition at the FAMA stage (Figure 4F-G and 5F-G). Based on these phenotypes, we propose that loop B mediates interactions with different scaffolds (Figure 9). A signal-promoting scaffold binds loop 7B and enforces MKK7 interactions with its cognate MKKK and MPK (Figure 9A). Such a scaffold would also explain why *in vivo* activity of MKK7^ED^ was always stronger than that of MKK5^DD^, even though *in vitro* assays showed an opposite pattern. On the other hand, we predict that a distinct scaffold recruited by loop 5B prevents MKK5^DD^ from inhibiting stomatal development at the FAMA stage (inhibitory scaffold) (Figure 9B). This prediction could also partially explain the behavior of certain chimeras. For example, the inhibitory scaffold would bind MKK7^ED^-5B through loop 5B and dampen MKK7^ED^ activities. Likewise, a signal-promoting scaffold would bind MKK5^DD^-7B through loop 7B, releasing the inhibition of MKK5^DD^.

**Figure 9:**
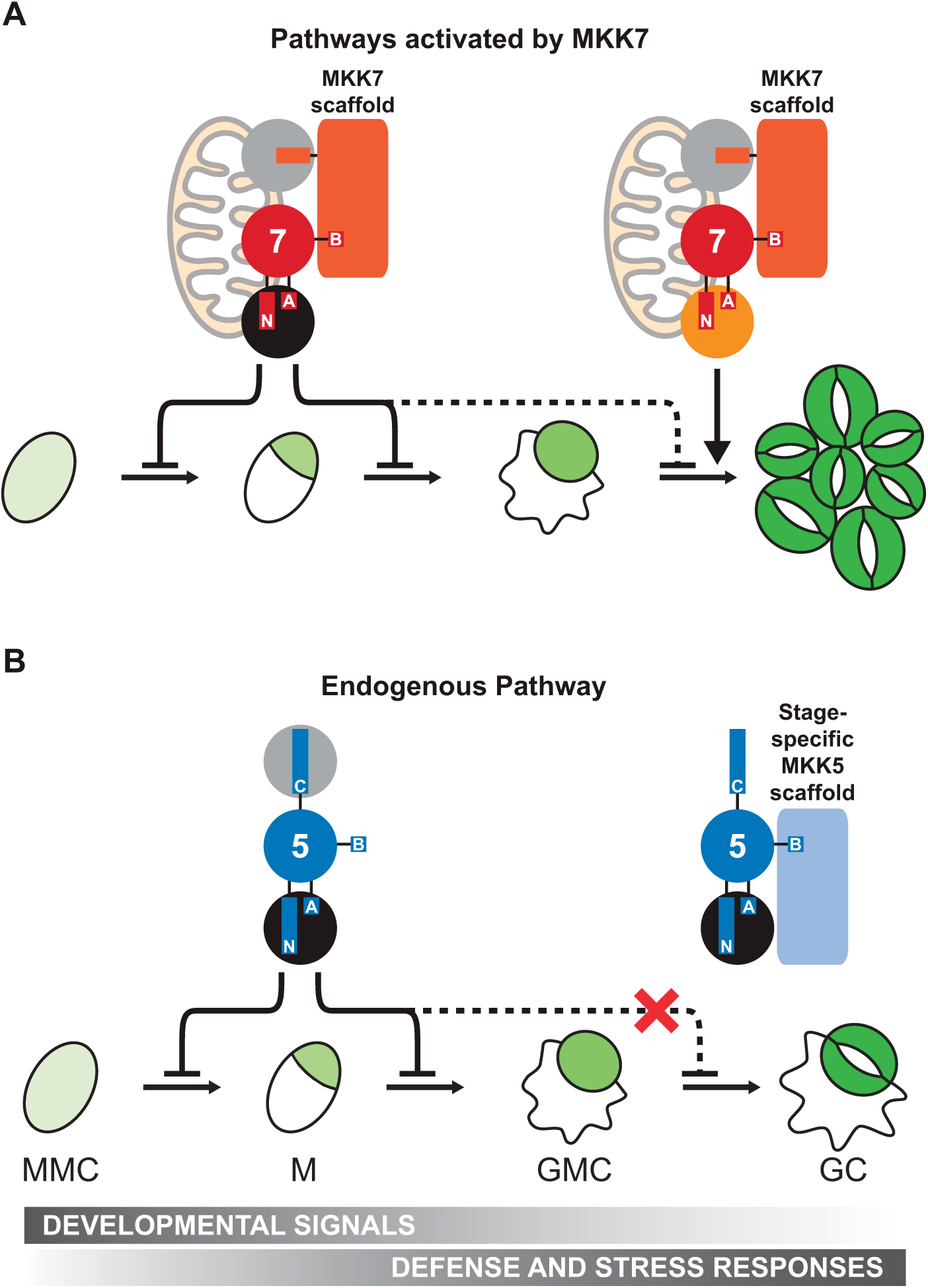
A new model derived from activities during stomatal development for the endogenous MKK5 pathway and another activated by MKK7. (A) In the MKK7 pathway, MKK7 mimics MKK5 in early development, forcing precursors to exit the stomatal lineage. At the last stage, MKK7 induces two phenotypes. Firstly, MKK7 induces stomatal clustering by means of an unknown MKK7-specific MPK (orange circle). This proliferation depends on MKK7 mitochondrial localization (Lampard et al., 2014). Secondly, MKK7 forces precursors to exit the lineage by escaping MKK5-specific regulation. An MKK7 scaffold enforces interactions between MKK7 and other components (MPKs and YODA, gray circles) within the network. (B) In the endogenous pathway, MKK5 and MPK3/6 (black circles) are involved in transducing developmental signals that inhibit stomatal lineage initiation and GMC commitment, and initiating defense and stress responses. At the last stage of development, a stage-specific MKK5 scaffold prevents MKK5 (and possibly MPK3/6) from inhibiting stomatal formation, only allowing defense and stress responses.

The existence of signal-enhancing and inhibiting MAPK scaffolds in plants is supported by recent findings (Zhao et al., 2014; Cheng et al., 2015; Zhang et al., 2015). While the relevance of the first type is quite intuitive, the second type is more controversial. The inhibitory scaffold EDR1 was identified in the context of pathogen defense. Current hypotheses are that EDR1 provides a failsafe against inadvertently activating defense or cell death programs when other (not pathogen-induced) cues activate MAPK signaling. EDR modulates MPK3 activity indirectly by interacting with MKK4/5 and regulating their abundance (Zhao et al., 2014). Interestingly, we observed in FAMA stage cells that MPK6 was only inhibited when the upstream MKK was also inhibited. The clearest evidence that MKK5-MPK3/6 are scaffolded in FAMA stage cells would be, of course, to identify the scaffold. We tested whether EDR1 worked to inhibit FAMA-stage expressed MKK5^DD^, however we failed to see a loss of inhibition in *edr1* (none seen in 20 independent T1s). EDR1 is a member of the large Raf-like MKKKs family (Ichimura et al., 2002) and potentially any of the 48 members of this family (alone or in combination) could serve as a stomatal scaffold. As testing the entire family in not technically feasible, we used stomatal lineage expression data (Adrian et al., 2015) to identify two close homologues of EDR1 expressed at the FAMA stage, MAP3Kδ1 (At1g11850) and MAP3Kδ5 (At4g24480). *MAP3Kδ1* and *MAP3Kδ5* mutants, however, were indistinguishable from WT in their response to FAMA stage MKK5^DD^ expression (0/20 independent T1s for each). Thus, the molecular identity of the stomatal scaffold remains of great future interest.

MKK4/5 face the problem of being used in early stages to repress stomatal progression, but being required for physiological regulation in guard cells. At the FAMA stage, MKK4/5 must therefore be actively rerouted from their previous role inhibiting stomatal development to allow terminal differentiation of guard cells (Figure 9). A negative scaffold acting late in the stomatal lineage to redirect MKK4/5 would provide an elegant solution to a complex signal integration problem.

## Supplemental Figure legends

**Supplemental Figure 1: Yeast two-hybrid assay with native and chimeric MKKs and MPK3/MPK6.** Representative yeast two-hybrid assay between MKK chimeras and MPK3/6 at 3 days of growth on control (-LW) and interaction (-LWH); three patches are serial (10 fold) dilutions.

**Supplemental Figure 2: *In vitro* kinase assays of MKKs and chimeras using kinase inactive (KI) MPK3 and MPK6 as substrates.** Phosphorylation assays were performed as described in methods. Samples were separated in SDS-PAGE, transferred to PVDF membranes and probed with anti-pERK antibody. Western blots were quantified and average of experiments was used as an estimation of *in vitro* activity. Each lane is labeled with the figure where the data is presented (grey bar marked *Figures*), the kinase used with a graphic representation and with its full name in bottom panel. Samples unassigned to any figure (labeled as “-” in *Figure*) were not presented in the main figures of this manuscript for the sake of brevity, but are included here to allow us to keep the blots intact. Red squares highlight samples that were not considered for quantification due to detection artifacts.

**Supplemental Figure 3: Multiple sequence alignment (Sievers et al., 2011) of catalytic domains in MPK kinases from mouse, rat, human and *Arabidopsis thaliana*.** A, Conserved subdomains (**Subdom.**) and consensus sequences (**Consen.**) are represented on top of the alignment and follow the same codes and convention for an alignment of 60 different kinases by Hanks and Hunter (Hanks and Hunter, 1995). In the consensus line: *uppercase letters,* invariant residues; *lowercase residues,* nearly invariant residues; o, positions conserving nonpolar residues; *, positions conserving polar residues; +, positions conserving small residues with near neutral polarity. Mammalian representative kinases selected for the alignment have been crystalized and belong to **AGC group** [cAMP-dependent kinase, cGMP-dependent kinase, etc.], **CAMK group** [Calcium-calmoduline-dependent protein kinase], **CMGC group** [cyclin-dependent kinase, mitogen-activated kinase, glycogen synthase kinase and cyclin-dependent-like kinase] and **STE group** [homologues of STE11 and STE20]. Gray boxes show CMGC insert (Goldsmith et al., 2007) in ERK2, CDK2 and p38, and Pro-rich sequence (PRS, involved in binding the scaffold MP1) in MEK1 and MEK2 were included in the alignment and cause an expansion of subdomain X. Secondary structure (**2° str.**) information is overlaid in the alignment (red for α-helices and yellow for β-strands). Conserved α- helices and β-strands are labeled following convention (Hanks and Hunter, 1995; Knight et al., 2007). Due to CMGC insert and PRS, αG helix is located in two different regions of the alignment and was named differently (residues underlined): **αG1** for MmPKA, HsARK-1, RnCaMKI, RnERK2, HsCDK2 and Hsp38; and **αG2** for HsMEK1/2 and AtMKK5/7. In green text, missing residues in crystal structures from HsMEK1 and HsMEK2 which include the PRS. B, Table provides general information and structural model names for kinases used in this comparison. C, Multiple sequence alignment of N- and C-termini from human MEK1/2 and Arabidopsis MKK5/7. Sequences highlighted in gray correspond to first and last three amino acids of the catalytic domains shown in A.

**Supplemental Figure 4: Multiple sequence alignment of C-termini in Arabidopsis and human MKKs.** Top panel, partial sequences for all Arabidopsis and human MKKs were aligned with ClustalOmega. Loops A and B defined for Arabidopsis MKKs are highlighted in gray. Number of first amino acid of the partial sequence is noted to the left of each sequence. The long C-terminal extension of AtMKK3 is truncated in this figure. Bottom panel, genes used in the alignment. Name of genes for Arabidopsis correspond to Arabidopsis Genome Initiative (AGI) codes and GeneCards (GC) for humans. NCBI GI, National Center for Biotechnology Information protein sequence identifier.

**Supplemental Figure 5: Multiple sequence alignment for MKK4, MKK5, MKK7 and MKK9.** Domains described in (Lampard et al., 2014) are highlighted in blue and green.

**Supplemental Figure 6: Representative phenotype and subcellular localization of transgenic seedlings expressing FAMAp:N7-MKK6^EE^-C7.**

**Supplemental Figure 7: *In vitro* kinase activity of MKKs does not correlate with their *in vivo* activity.** Linear regressions for MPK3 or MPK6 activity versus SPCH stage (A) or FAMA stage (B) activities, with their formulas and R^2^, are displayed in the figure. FAMA stage activity was calculated as the addition of phenotypes different than normal (Inhibited, Small and Large clusters in Table 1).

